# Genome-wide Analysis of Differential Transcriptional and Epigenetic Variability Across Human Immune Cell Types

**DOI:** 10.1101/083246

**Authors:** Simone Ecker, Lu Chen, Vera Pancaldi, Frederik O. Bagger, José Maria Fernandez, Enrique Carrillo de Santa Pau, David Juan, Alice L. Mann, Stephen Watt, Francesco Paolo Casale, Nikos Sidiropoulos, Nicolas Rapin, Angelika Merkel, BLUEPRINT Consortium, Henk Stunnenberg, Oliver Stegle, Mattia Frontini, Kate Downes, Tomi Pastinen, Taco W. Kuijpers, Daniel Rico, Alfonso Valencia, Stephan Beck, Nicole Soranzo, Dirk S. Paul

**Affiliations:** Structural Biology and Biocomputing Programme, Spanish National Cancer Research Center (CNIO), Melchor Fernández Almagro 3, 28029 Madrid, Spain; UCL Cancer Institute, University College London, 72 Huntley Street, London, WC1E 6BT, UK; Department of Human Genetics, Wellcome Trust Sanger Institute, Wellcome Trust Genome Campus, Hinxton, Cambridge, CB10 1HH, UK; Department of Haematology, University of Cambridge, Cambridge Biomedical Campus, Long Road, Cambridge, CB2 0PT, UK; National Health Service (NHS) Blood and Transplant, Cambridge Biomedical Campus, Long Road, Cambridge, CB2 0PT, UK; European Molecular Biology Laboratory, European Bioinformatics Institute, Wellcome Trust Genome Campus, Hinxton, Cambridge, CB10 1SD, UK; The Finsen Laboratory, Rigshospitalet, Faculty of Health Sciences, University of Copenhagen, Ole Maaløes Vej 5, 2200, Copenhagen, Denmark; Biotech Research and Innovation Centre (BRIC), University of Copenhagen, Ole Maaløes Vej 5, 2200, Copenhagen, Denmark; The Bioinformatics Centre, Department of Biology, Faculty of Natural Sciences, University of Copenhagen, Ole Maaløes Vej 5, 2200, Copenhagen, Denmark; National Center for Genomic Analysis (CNAG), Center for Genomic Regulation (CRG), Barcelona Institute of Science and Technology, Carrer Baldiri i Reixac 4, 08028 Barcelona, Spain; Department of Molecular Biology, Radboud University, Faculty of Science, Nijmegen, 6525 GA, The Netherlands; British Heart Foundation Centre of Excellence, University of Cambridge, Cambridge Biomedical Campus, Long Road, Cambridge, CB2 0PT, UK; Department of Human Genetics, McGill University, 740 Dr. Penfield, Montreal, H3A 0G1, Canada; Blood Cell Research, Sanquin Research and Landsteiner Laboratory, Plesmanlaan 125, Amsterdam, 1066CX, The Netherlands; Emma Children’s Hospital, Academic Medical Center (AMC), University of Amsterdam, Location H7-230, Meibergdreef 9, Amsterdam, 1105AX, The Netherlands; Cardiovascular Epidemiology Unit, Department of Public Health and Primary Care, University of Cambridge, Strangeways Research Laboratory, Wort’s Causeway, Cambridge, CB1 8RN, UK

**Keywords:** Differential variability, phenotypic plasticity, heterogeneity, immune cells, monocytes, neutrophils, T cells, gene expression, DNA methylation

## Abstract

**Background:** A healthy immune system requires immune cells that adapt rapidly to environmental challenges. This phenotypic plasticity can be mediated by transcriptional and epigenetic variability.

**Results:** We applied a novel analytical approach to measure and compare transcriptional and epigenetic variability genome-wide across CD14^+^CD16^−^ monocytes, CD66b^+^CD16^+^ neutrophils, and CD4^+^CD45RA^+^ naïve T cells, from the same 125 healthy individuals. We discovered substantially increased variability in neutrophils compared to monocytes and T cells. In neutrophils, genes with hypervariable expression were found to be implicated in key immune pathways and to associate with cellular properties and environmental exposure. We also observed increased sex-specific gene expression differences in neutrophils. Neutrophil-specific DNA methylation hypervariable sites were enriched at dynamic chromatin regions and active enhancers.

**Conclusions:** Our data highlight the importance of transcriptional and epigenetic variability for the neutrophils’ key role as the first responders to inflammatory stimuli. We provide a resource to enable further functional studies into the plasticity of immune cells, which can be accessed from: http://blueprint-dev.bioinfo.cnio.es/WP10/hypervariability.

## Background

Phenotypic plasticity is fundamental to human immunity, allowing rapid cellular adaptation in response to changing environmental conditions [1]. Plasticity of immune cells can be influenced by the variability of cellular traits including gene expression and DNA methylation. The stochastic nature inherent to cellular processes such as gene regulation gives rise to cell-to-cell variation, enhancing survival under adverse conditions and stress [2–4]. Environmental stimuli including temperature, hormone levels, and invading pathogens further affect the expression of genes in a tissue- and temporal-dependent fashion [2,4,5].

Rapid and effective response to a stimulus is facilitated and intensified if the cellular trait already exhibits large stochastic fluctuations in the absence of the stimulus [3]. For example, while genes involved in stress response tend to be highly variable [3,6,7], genes involved in essential cellular functions, such as protein synthesis and metabolism, demonstrate less variable expression levels [8,9]. B and T cells utilize genetic recombination to generate a highly diverse repertoire of immunoglobulins and T cell surface receptors, respectively. In addition, immune responses are driven by the variability of key signaling molecules and transcription factors not controlled by genetic factors [10,11]. Epigenetic states, including DNA methylation, also contribute to plastic gene expression during cell fate commitment, thus enhancing fitness in response to external cues [12,13].

Transcriptional and epigenetic heterogeneity that is measured across individuals emerges from different origins. While intra-individual variability can relate to different cellular properties in response to external signals, such as cell activation and communication [3,7,14], inter-individual variability can relate to differences between the individuals, including genetic makeup, age, sex, and lifestyle. Importantly, it has also been demonstrated that inter-individual variability can serve as an appropriate proxy for intra-individual variability at the level of single cells [7,14,15]. Both transcriptional and epigenetic variability have been shown to strongly correlate with the development and progression of human diseases [12,16,17]. For example, gene expression variability has been linked to human immunodeficiency virus (HIV) susceptibility [18], neurological disorders [18,19], and cancer [20,21]. Hypervariable DNA methylation loci can be used as biomarkers to predict the risk of neoplastic transformation in stages prior to neoplasia [22,23].

The extent and functional interpretation of transcriptional and epigenetic variability has not been systematically investigated genome-wide across multiple immune cell types in the general population. Here, we applied a novel analytical approach to measure differential variability of gene expression and DNA methylation in three major immune cell types: CD14^+^CD16^−^ classical monocytes, CD66b^+^CD16^+^ neutrophils, and CD4^+^CD45RA^+^ ‘phenotypically naïve’ T cells. This matched panel of cell types was derived from the same 125 healthy individuals. We show that neutrophils exhibit substantially increased variability of both gene expression and DNA methylation patterns, compared to monocytes and T cells, consistent with these cells’ key role as the first line of host defense. We annotated hypervariable genes and CpGs to known homeostatic and pathogenic immune processes, and further found subsets of genes correlating with genetic makeup, donor demographic and lifestyle factors. Our data further reveal potential molecular mechanisms of immune responses to environmental stimuli, and provide a resource to enable future functional studies into the phenotypic plasticity of human immune cells in health and disease.

## Results

### Deep molecular profiling of immune cells in the BLUEPRINT Human Variation Panel

The analyses described in this study are based on the publicly available resource provided by the BLUEPRINT Human Variation Panel [24]. The resource contains genome-wide molecular profiles of CD14^+^CD16^−^ classical monocytes, CD66bn^+^CD16^+^ neutrophils, and CD4^+^CD45RA^+^ naïve T cells. These leukocyte types were chosen due to their important role in mediating immune cell processes, their relative abundance in peripheral blood allowing for examination of multiple cellular traits, as well as the availability of experimental protocols to prepare cell populations of high purity (>95%). Monocytes and neutrophils are myeloid cells that share the same bone-marrow residing granulocyte-macrophage precursor cell. Monocytes migrate to sites of infection and differentiate into macrophages and dendritic cells to induce an immune response. As part of the innate immune system, neutrophils move within minutes to sites of infection during the acute phase of inflammation. Naïve T cells are lymphoid cells that are part of the adaptive immune system, representing mature helper T cells that have not yet recognized their cognate antigen.

Across an initial cohort of 202 healthy individuals representative of the UK population, purified preparations of these primary cells were probed for gene expression using total RNA sequencing (RNA-seq) and DNA methylation using Illumina Infinium HumanMethylation450 BeadChips (‘450K arrays’). Detailed information about the experimental and analytical strategies for quantifying these cellular traits are provided in the Methods section. Figures S1 and S2 give an overview of the data quality assessment of the gene expression and DNA methylation data sets, respectively. All individuals were further profiled for DNA sequence variation using whole-genome sequencing to allow for cell type-dependent, quantitative assessment of the genetic and epigenetic determinants of transcriptional variance [24].

In this study, we exploited this resource, selecting all 125 donors for whom matched gene expression and DNA methylation data sets were available across the three immune cell types. The key analytical advance of the work presented here concerns the measurement and interpretation of differential variability. That is, the identification of loci at which gene expression and DNA methylation levels show significantly greater variation within one cell type as compared to the other cell types. An overview of the study design and analytical concept is provided in Figure 1A.

**Figure 1.**
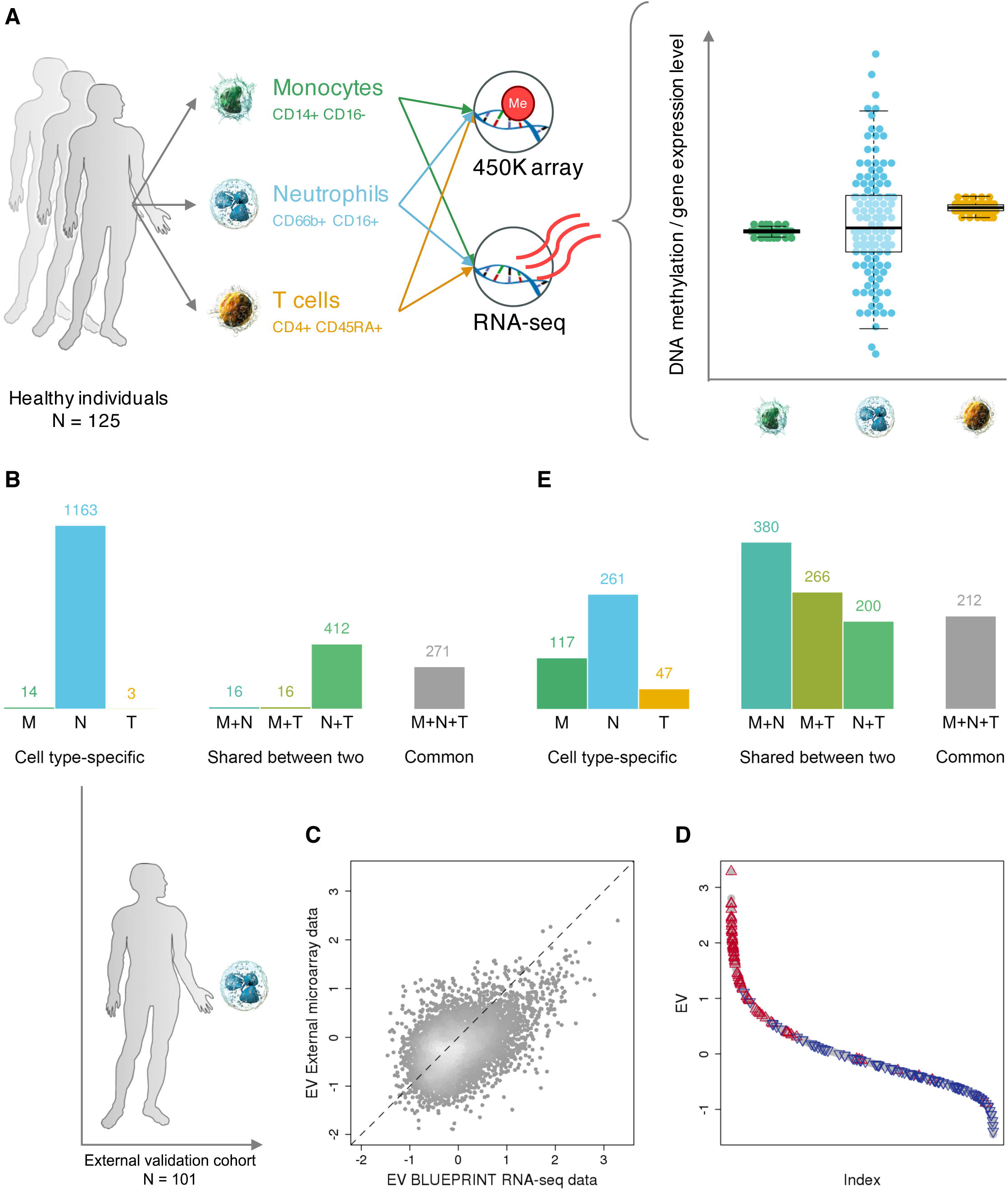
Differential variability of gene expression and DNA methylation across three immune cell types. (A) Study design and analytical approach. Hypervariable genes and CpGs were identified using a (A) combined statistical approach at stringent significance thresholds, i.e. Benjamini-Hochberg (BH)- corrected *P* <0.05 and gene expression/DNA methylation variability measurement (EV/MV) difference ≥10% relative to the observed range. (B) Barplots showing the number of statistically significant hypervariable genes (HVGs) that are (A) cell type-specific, shared between two, or common to all three of the studied immune cell types. (C) Scatter plot of the EV-values of 6,138 genes assessed in our data set versus the replication set. (A) We found good concordance between the two independent cohorts, despite the application of different analytical platforms (Pearson’s r = 0.48, *P* <2.2×10-16). (D) Ranking of all 11,980 protein-coding genes analyzed in our study according to EV-values (i.e. (A) from high to low EV-values). We highlight the 100 genes that showed the highest and lowest EV-values in the independent replication data set in red and blue, respectively. (E)) Barplots showing the number of hypervariable CpG positions (HVPs). (A) *Abbreviations: M=monocytes, N=neutrophils, T=naïve T cells.*

### Genome-wide patterns of differential gene expression variability across immune cell types

We first assessed inter-individual expression variability of 11,980 protein-coding, autosomal genes that showed robust expression in monocytes, neutrophils, and T cells (Methods). We applied an improved analytical approach for the assessment of differential variability (Methods), taking into account the strong negative correlation between mean gene expression levels and expression variability (Figure S3).

Figure 1B gives an overview of the number of identified hypervariable genes (HVGs) that are cell type-specific, shared between two, or common to all three of the studied immune cell types. Neutrophils were found to have the largest number of HVGs overall (n=1,862), as well as of cell type-specific HVGs (n=1,163). In contrast, we found only a small number of cell type-specific HVGs in monocytes and T cells (n=14 and n=3, respectively). In addition, we identified 271 genes that were highly variable across all three immune cell types using a rank-based approach (Methods). Mature neutrophils (as profiled here) show low proliferative capacity and reduced transcriptional and translational activity [25,26]. The latter could potentially impede comparable assessment of differential variability, if the relationship between variability and mean expression levels was not taken into account. Thus, using our analytical approach, we assessed and confirmed that overall reduced gene expression levels did not technically confound the observed increased variability of gene expression levels in neutrophils (Figure S3).

We then aimed to replicate the detected hypervariable gene levels in an independent sample cohort. We retrieved a gene expression data set generated using Illumina Human HT-12 v4 Expression BeadChips consisting of CD16^+^ neutrophils derived from 101 healthy individuals [27]. Of the 11,023 gene probes assessed on the array platform, 6,138 could be assigned to a corresponding gene identifier in our data set. Neutrophil-specific hypervariable genes measured using RNA-seq were also found to be hypervariable using expression arrays in the independent cohort of healthy individuals (Figures 1C and 1D).

In summary, we devised and assessed a novel method for the identification of differential gene expression variability. Overall, we found strongly increased variability of gene expression in neutrophils compared to monocytes and T cells, and replicated the detected neutrophil-specific hypervariable gene patterns in an external cohort.

### Biological significance of differentially variable genes across immune cell types

Next, we explored the characteristics of the identified hypervariable genes. We performed ontology enrichment analysis of gene sets using the GOseq algorithm [28]. This method takes into account the effect of selection bias in RNA-seq data that can arise due to gene length differences [28]. Table S1 and Table S2 summarize the annotation data of all identified HVGs and observed gene ontology enrichment patterns, respectively.

Genes showing expression hypervariability across all three cell types were enriched in biological processes related to chemotaxis, migration, and exocytosis (Table S2). For neutrophil-specific HVGs, we found gene ontology enrichment in oxidoreductase activity and cellular processes related to virus response and parasitism (Table S2). Notable genes among those with hypervariable expression values were *CD9* (Figure 2A), *CAPN2* (Figure 2B), and *FYN* (Figure C). *CD9* showed increased variability across all three cell types. The gene encodes the CD9 antigen, a member of the tetraspanin family. It functions as cell surface protein that forms complexes with integrins to modulate cell adhesion and migration and mediate signal transduction [29,30]. The neutrophil-specific HVGs *CAPN2* and *FYN* encode a calcium-activated neutral protease involved in neutrophil chemotaxis [31] and a tyrosine-protein kinase implicated in intracellular signal transduction [32], respectively.

**Figure 2.**
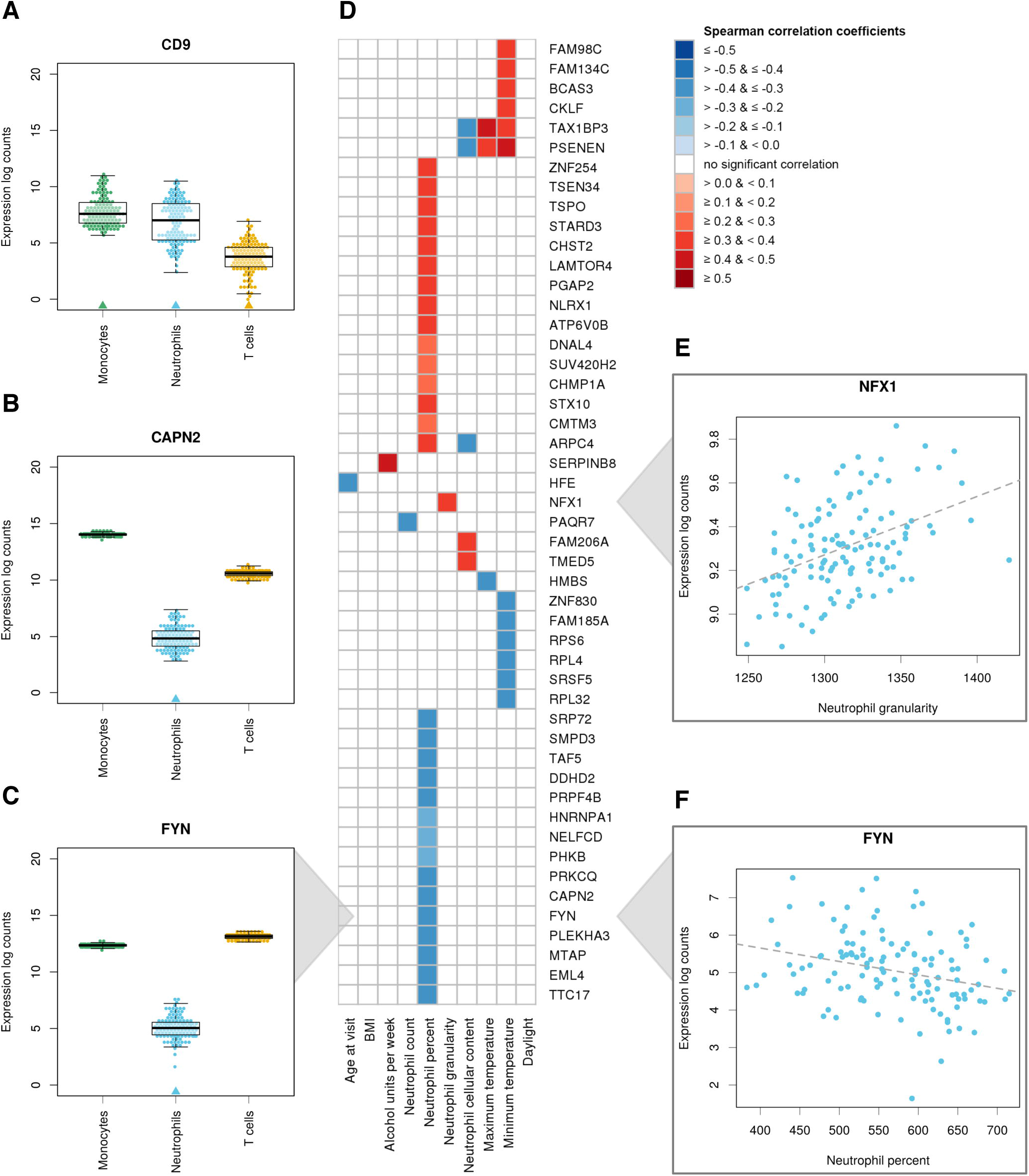
Characterization of cell type-specific hypervariable genes. (A–C) Increased expression variability of the genes *CD9*, *CAPN2*, and *FYN* across three immune cell types. For each cell type, data points represent the expression values of the indicated gene in one individual. Cell types marked by an arrow were found to show significantly increased variability compared to the other two cell types. While *CD9* was found to be hypervariable in all three cell types, *CAPN2* and *FYN* show increased variability only in neutrophils, if contrasted to monocytes and T cells. (D) Heatmap of Spearman’s correlation coefficients showing neutrophil-specific HVGs that (D) associated with various donor-specific quantitative traits. A total of 49 genes with increased inter-individual variability showed a significant association with at least one of the measured traits (BH-corrected *P* <0.05, Spearman’s rank correlation). (E)) Scatter plot of *NFX1* gene expression levels versus neutrophil granularity. (D) (F)) Scatter plot of *FYN* gene expression levels versus neutrophil percentage. (D)

Taken together, functional enrichment of hypervariable gene sets revealed that many of the identified hypervariable genes are involved in mediating immune-related processes. This suggests that neutrophils exhibit specific gene loci that are highly adaptable to external cues.

### Determinants of inter-individual cell type-specific gene expression variability

Following the discovery and characterization of genes that present hypervariable expression levels between individuals, we next aimed to delineate potential sources of heterogeneity that can be associated to differences between individuals. We hypothesized that these sources mainly relate to genetic variation, age, sex, and lifestyle factors.

First, we determined the subset of cell type-specific HVGs that correlated with genetic variants. We retrieved gene sets with a local (*cis*) genetic component designated by expression quantitative trait locus (eQTL) and variance decomposition analyses, as described in the BLUEPRINT Human Variation Panel (Figure S4A). In neutrophils, we found that 638 of the 1,163 cell-specific HVGs (55%) associate with *cis* genetic variants (Table S1), at least partly explaining the observed gene expression variability. These data are consistent with previous reports, highlighting the role of genetic variants in mediating transcriptional variance [33–35].

Second, we correlated cell type-specific HVGs with various quantitative traits measured in individual donors: demographic information (age, body mass index, and alcohol consumption); cellular parameters as assessed by a Sysmex hematology analyzer (e.g. cell count and size); and season (i.e. minimum/maximum temperature and daylight hours of the day on which blood was drawn). The results of this analysis are provided in Tables S1 and S3. In neutrophils, we identified HVGs that show significant association with at least one of the measured traits (Figure 2D). For example, we found *NFX1*, a nuclear transcription factor that regulates *HLA-DRA* gene transcription [36], to associate with neutrophil granularity (Figure 2E). An increase in neutrophil granularity can be reflective of a potential infection; this parameter is routinely monitored in a clinical setting. *FYN* gene levels (reported above) were negatively correlated with neutrophil percentage (Figure 2F).

Third, we investigated whether sex was an important source of inter-individual (autosomal) gene expression variability. We found only two of the 1,163 neutrophil-specific HVGs to be differentially expressed between sexes, *SEPT4* and *TMEM63C* (Figure S5A), and high expression variability was observed for both sexes in these genes. However, in neutrophils we identified a surprisingly large number of sex-specific differentially expressed genes of small effect size, which corresponded to important immune cell functions. We present a detailed analysis of these genes below.

In conclusion, we found that genetic makeup was an important determinant of transcriptional variability. Donor demographic and lifestyle factors also contributed towards transcriptional variability.

### Neutrophil-specific hypervariable genes not mediated by *cis* genetic effects

Next, we studied in detail the subset of neutrophil-specific genes that showed hypervariable expression but did not associate with local genetic variants (n=525). Although some of these genes could be mediated by distal (*trans*) genetic factors not detected in the BLUEPRINT Human Variation Panel, it is conceivable that expression heterogeneity of this gene set was primarily due to external triggers or stochastic fluctuations.

We generated a correlation matrix of expression levels of the 525 HVGs, and identified clusters of correlated genes that may act in concert or be co-regulated. The identified co-expression network contained 259 connected genes, and consisted of three distinct gene modules (Figure 3). We inferred biological functions corresponding to the three gene modules. All modules were highly enriched for genes with important immune-related functions.

**Figure 3.**
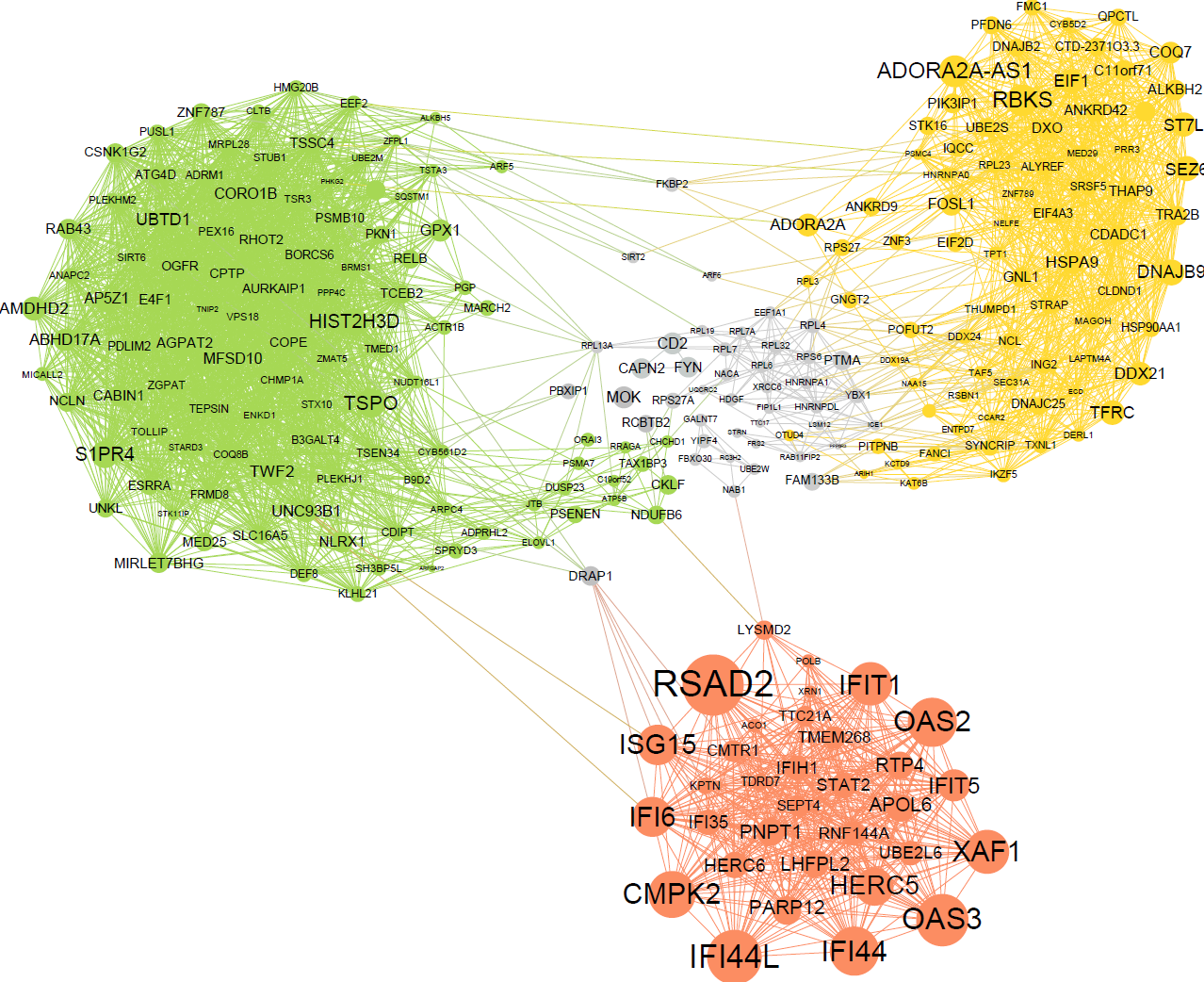
Gene network and pathway analysis of neutrophil-specific HVGs not mediated by *cis* genetic effects. Co-expression network of neutrophil-specific HVGs that did not correlate with genetic variants in *cis*, as reported in the BLUEPRINT Human Variation Panel. We identified three gene modules, shown in green, yellow, and red. These modules were highly enriched for important biological functions in immune cells (Table S4). Nodes represent genes, and edges represent correlations in these genes’ expression values. Node sizes are determined by expression variability of the corresponding gene, with bigger nodes indicating higher EV-values. Nodes colored in gray belong to several smaller gene clusters connecting the three main clusters of the network.

The first and largest gene module (n=105 genes, green color) showed enrichment for inclusion body, receptor signaling, and immune response activation. The second module (n=78 genes, yellow color) was enriched in biological processes related to RNA processing and chaperone binding. The third gene module (n=33 genes, red color), contained many genes with particularly high variation in their expression patterns. *RSAD2*, an interferon-inducible antiviral protein, showed highest variability, among many other interferon-inducible genes present in module three. These genes are essential in innate immune response to viral infections [37]. Gene ontology and pathway analyses of all genes in the network module further showed a strong enrichment for response to type I interferon, and several viral disease pathways including influenza A, herpes simplex, and hepatitis (Figure S6). A detailed functional annotation of all three network modules is provided in Table S4.

### Sex-specific differential gene expression across immune cell types

In our analysis, we did not detect differences in mean gene expression levels between male and female donors with log-fold change ≥1 – albeit with two exceptions in neutrophils (Figure S5A). Nonetheless, when no minimum log-fold change criterion was applied, we found that sex-dependent mean expression of autosomal genes (Figure S5B) was highly abundant in neutrophils (n=3,357 genes), compared to T cells (n=895) and monocytes (n=64).

As many autoimmune diseases have a higher incidence in females, and females show generally elevated immune responses compared to males [38], we hypothesized that genes with elevated gene expression levels in females may account for the increased incidence rates. Indeed, genes with higher mean expression levels in neutrophils derived from females (n=682) were enriched in immune response and related pathways (Table S5). In contrast, genes with increased mean expression in male donors (n=2,675) were enriched in basic cellular processes, such as RNA processing and translation (Table S5). In addition, in male donors, genes were strongly enriched in cellular compartments, such as nuclear lumen (Table S5).

### Genome-wide patterns of differential DNA methylation variability across immune cell types

Following the analyses of differential gene expression variability, we then applied our improved analytical approach to determine inter-individual variability of DNA methylation levels at 440,905 CpG sites (Methods). Again, our method accounted for confounding effects due to the correlation between mean and variability measurements (Figure S7).

Concordant with our findings for gene expression variability (Figure 1B), we found that neutrophils had the largest number of hypervariable CpG positions (HVPs) overall (n=1,053), as well as cell-specific HVPs (n=261). Neutrophils and monocytes shared a considerable number of HVPs (n=380) in contrast to T cells (Figure 1E). Finally, we identified 212 HVPs common to all three cell types. An overview of the number of HVPs is shown in Figure 1E.

Following the discovery of HVPs, we examined whether these sites were overrepresented at particular gene elements and epigenomic features. To this end, we focused on cell type-specific HVPs, correlating their DNA methylation levels with distinct cellular characteristics and molecular pathways. In Table S6, we summarize the detailed annotation of all HVPs across the three profiled immune cell types. In neutrophils, we found that cell type-specific HVPs were depleted at CpG islands, which typically occur near transcription start sites (*P* = 6.37×10-19, hypergeometric test; Figure 4A), and enriched at intergenic regions (*P* = 0.03; Figure 4B).

**Figure 4.**
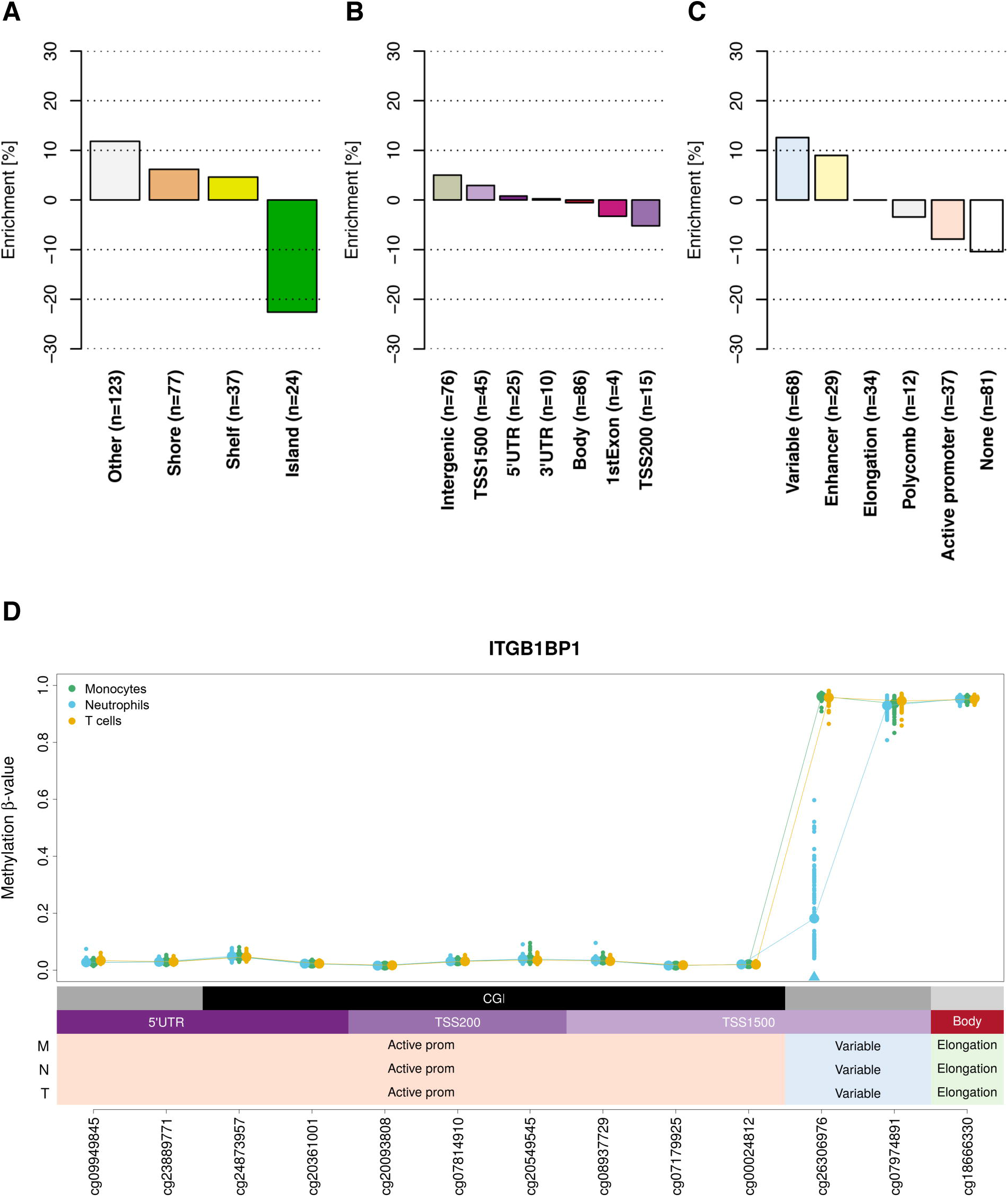
Functional annotation of neutrophil-specific hypervariable CpG positions. (A) Enrichment of neutrophil-specific HVPs (n=261) at genomic features. We found neutrophil- (A) specific HVPs to be depleted at CpG islands (*P* = 6.37×10-19, hypergeometric test). (B) Enrichment of neutrophil-specific HVPs at gene elements. Neutrophil-specific HVPs were (A) enriched at intergenic regions (*P* = 0.03). (C) Enrichment of neutrophil-specific HVPs at distinct reference chromatin states in neutrophils. (A) The HVPs were enriched at enhancer (*P* = 1.32×10-12) and ‘variable’ (*P* = 3.81×10-8) chromatin states. A ‘variable’ chromatin state denotes a state that was observed in less than 80% of the biological replicates (n ≥5) within a given cell type, and indicates dynamic changes of local chromatin structure. (D) Regional plot of an exemplar neutrophil-specific HVP mapping to the promoter of the (A) *ITGB1BP1* gene, encoding the integrin beta 1 binding protein 1. The statistically significant HVP is indicated with an arrow. For each cell type, data points represent the DNA methylation β-values (y-axis) at the indicated CpGs (x-axis) in one individual. For each CpG site, we calculated the mean DNA methylation value (indicated with a larger data point). Every CpG site is annotated with regards to genomic feature, gene element, and chromatin state. *Abbreviations: M=monocytes, N=neutrophils, T=naïve T cells, TSS=transcription start site, CGI=CpG island, UTR=untranslated region, prom=promoter.*

We hypothesized that cell type-specific HVPs localize at distal gene regulatory elements such as enhancer sequences, of which many are known to be also cell type-specific [39]. To test this hypothesis, we retrieved reference chromatin state maps of primary human monocytes, neutrophils, and T cells from the data repository provided by the BLUEPRINT Consortium [40]. Chromatin states are defined as spatially coherent and biologically meaningful combinations of multiple chromatin marks [41,42]. A total of five chromatin states were designated, which corresponded to functionally distinct genomic regions, namely active promoters, enhancers, and regions related to transcriptional elongation and polycomb-repression. In addition, a ‘variable’ chromatin state was defined here, indicating frequent changes of local chromatin structure across samples of the same cell type. Indeed, neutrophil-specific HVPs were found to be strongly enriched in the enhancer (*P* = 1.32×10-12, hypergeometric test; Figure 4C) and variable chromatin states (*P* = 3.81×10-8; Figure 4C).

### Biological significance of immune cell type-specific hypervariable CpGs

To interpret the potential cellular and biological implications of cell type-specific hypervariable CpGs, we annotated the genes in close proximity to each CpG using the Genomic Regions Enrichment of Annotations Tool (GREAT) [43]. This tool is valuable in assigning putative functions to sets of non-coding genomic regions [43].

Overall, we found enrichment in gene ontology terms attributed to genes close to HVPs in a cell type-dependent context (Table S7). For example, genes located near neutrophil-specific HVPs were enriched in gene signatures related to acute *Streptococcus pneumoniae* infection and cysteine synthase activity; the latter molecular process is important to hold off infections [44]. Consistent with established neutrophil function, this suggests that the identified HVPs play a role in regulating the expression of neutrophil-specific genes in response to infection.

In Figure 4D, we provide an example of a neutrophil-specific HVP at the promoter of the *ITGB1BP1* gene, encoding the integrin beta 1 binding protein 1. Integrins are essential cell adhesion proteins that induce intracellular signaling pathways upon activation by matrix binding [45,46]. They function as signal transducers allowing for rapid responses to cell surface signals [46]. Notably, the highlighted HVP mapped to a variable chromatin state at this locus, indicating that it influences local chromatin dynamics upon an internal or external trigger (Figure 4D).

In conclusion, we showed that cell type-specific HVPs clustered in enhancer and dynamic chromatin states at intergenic regions, suggesting they play a role in the regulation of cell type-specific gene expression programs in response to environmental changes. Genes in proximity to HVPs were enriched in gene sets relevant to important immunological functions.

### Determinants of inter-individual cell type-specific DNA methylation variability

Subsequent to the identification and annotation of CpGs with hypervariable DNA methylation levels, we explored potential reasons for the discovered inter-individual DNA methylation heterogeneity.

In agreement with our findings for gene expression variability, we determined that a large proportion of cell type-specific HVPs correlated with *cis* genetic variants reported in the BLUEPRINT Human Variation Panel (Figure S4B). In neutrophils, we found that 167 of the 261 cell type-specific HVPs (64%) associated with DNA methylation quantitative trait loci (Table S6). Our data further revealed that none of the cell type-specific HVPs were differentially methylated between male and female donors. The complete numerical results of all correlation analyses are provided in Table S8.

HVPs specific to monocytes showed frequent association with seasonal effects, such as temperature and daylight (n=12/117 HVPs; Figure S8). This finding is consistent with recent analyses reporting fluctuations of gene expression levels in monocytes depending on season and circadian rhythm [47]. Many CD4^+^ T cell-specific HVPs particularly correlated with donor age (n=14/46 HVPs; Figure S8), in line with previous findings on age-related DNA methylation changes in T cells [48,49]. These alterations are especially interesting in the context of immunosenescence, for which dysregulation in T cell function is thought to play a crucial role [50,51]. Naïve CD4^+^ T cells have further been reported to become progressively longer-lived with increasing age [52], which possibly also impacts their DNA methylation patterns.

### Correlation of DNA methylation variability with transcriptional output

DNA methylation at active gene elements can directly control the regulation of gene expression. While methylated gene promoters usually lead to transcriptional silencing, methylated gene bodies typically lead to transcriptional activation [53]. We next aimed to probe this paradigm in the context of gene expression and DNA methylation variability.

We measured the correlation of DNA methylation variability with transcriptional output at the level of single genes. Specifically, we studied cell type-specific HVPs that map to gene promoters and bodies, correlating their DNA methylation level with the gene expression level in the same individuals. At promoters, 30.1% (range, 23.5^−^33.3%) of HVPs showed a negative correlation with gene expression (Figure 5A), in support of the conventional role of DNA methylation in gene repression. At gene bodies, a small subset of HVPs (5.0%; range, 0.0^−^10.8%) showed a positive correlation with gene expression (Figure 5B). Table S9 gives a full account of these genes and numeric results.

**Figure 5.**
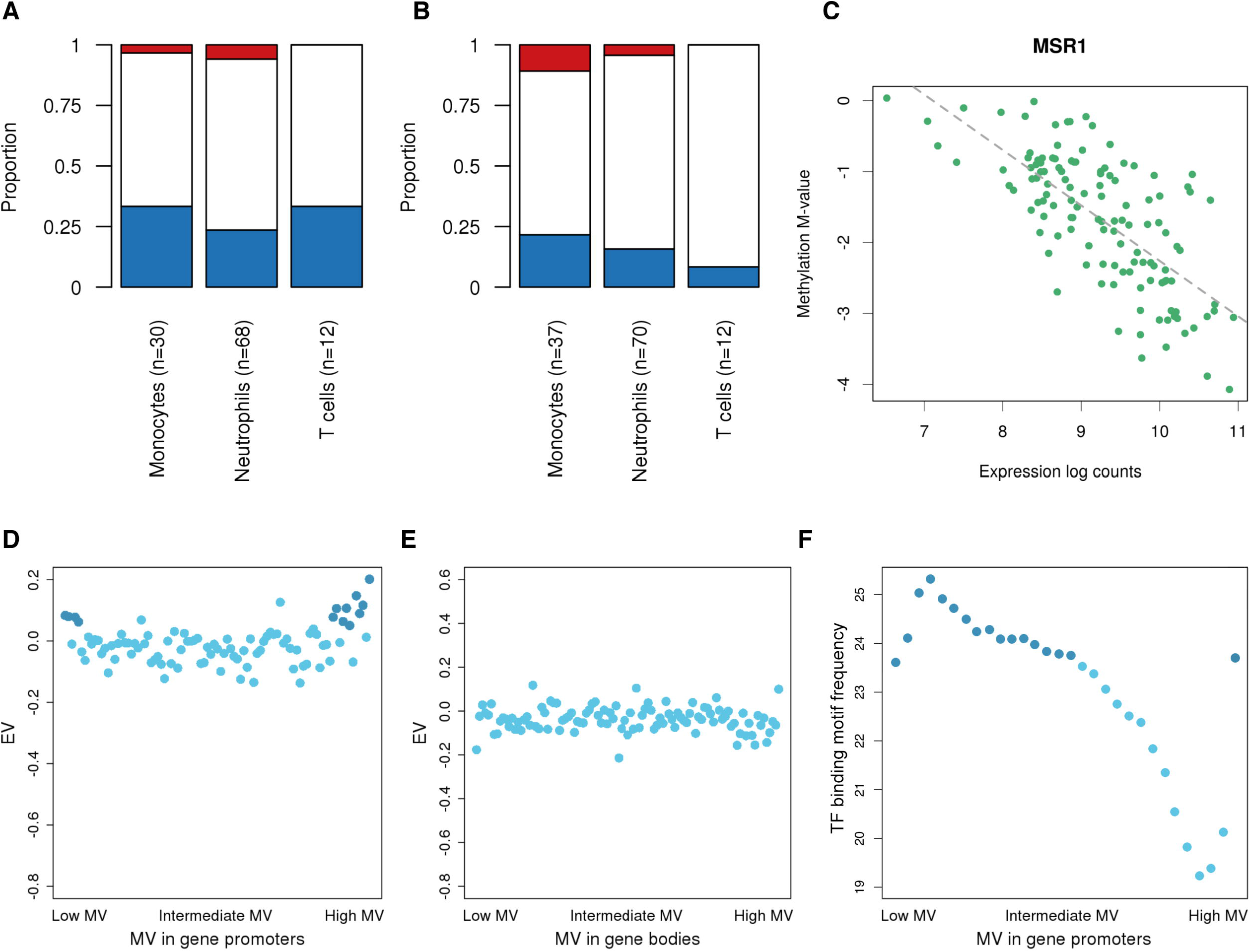
Relationship between DNA methylation and gene expression. (A) Barplots showing the proportion of cell type-specific HVPs that map to gene promoters and (A) are positively (red), negatively (blue), or not (white) associated with gene expression levels at BH-corrected *P* <0.05 (Spearman’s rank correlation). We found that around one third of these HVPs (30.1%; range 23.5^−^33.3%) is negatively correlated with gene expression. (B) Barplots as shown in panel (A) but for HVPs that map to gene bodies. (A) (C) Scatter plot showing the negative correlation of *MSR1* promoter DNA methylation with gene (A) expression in monocytes (r = −0.70, *P* <2.2×10-16; Spearman’s rank correlation). (D) Correlation between DNA methylation variability and gene expression variability at gene (A) promoters in neutrophils. First, gene-wise MV-values were calculated. Then, the values were ordered from low to high MV-value, grouped together in bins of 100 genes, and plotted against the EV-values, maintaining the ordering by MV-values. This binning strategy was applied to reduce the complexity of the data. HVPs at gene promoters were defined as CpG sites annotated to TSS1500, TSS200, 5’UTR, and first exon, according to the Illumina 450K array annotation manifest. Darker data points indicate the subset of bins that is further discussed in the Results section. (E) Same scatter plot as shown in panel (D) but for HVPs that map to gene bodies. HVPs at gene (A) bodies were defined as CpGs annotated to body and 3’UTR, according to the 450K array annotation manifest. (F) Scatter plot of the number of consensus transcription factor binding motifs at promoter regions (A) versus MV-values in neutrophils. Promoter regions were defined as ±500 bp around the transcription start site. Darker data points indicate the subset of bins that is further discussed in the Results section. *Abbreviations: MV=DNA methylation variability, TF=transcription factor.*

An example is provided in Figure 5C, showing a monocyte-specific HVP at the gene promoter of *MSR1*. At this CpG site, DNA methylation levels were significantly correlated with gene repression (BH-corrected *P* <2.2×10-16, Spearman’s rank correlation). *MSR1*, encoding the CD204 antigen, is involved in endocytosis of modified low-density lipoproteins.

### Relationship between DNA methylation variability and gene expression variability

Finally, we examined global patterns of DNA methylation variability in relation to transcriptional variability. In neutrophils, highly variable gene expression levels were observed at promoters exhibiting highly variable DNA methylation levels, and also at promoters showing very stable DNA methylation levels (Figure 5D). For DNA methylation variability at gene bodies, this relationship was weaker and showed a linear tendency (Figure 5E). Importantly, these global patterns were consistent across all three immune cell types (Figure S9).

To characterize these promoter regions further, we counted the number of transcription factor binding motifs at these regions (Methods). We found an accumulation of binding motifs at promoters presenting either highly variable or very stable DNA methylation levels (Figure 5F and S8). Next, we explored the properties of the 100 genes that showed both the highest expression variability and the highest DNA methylation variability at their promoters. We found that of the genes in each cell type, 66 were common to all three cell types; in turn, 10 of these 66 genes encode transcription factors. For example, in neutrophils this included *ELF1*, a transcriptional regulator of genes involved in immune response signaling pathways [54]. Neutrophil-specific HVGs were also enriched at genes with promoter sequences that contain the consensus binding motif of *ELF1* (BH-corrected *P* = 1.2×10-5; MSigDB analysis).

Taken together, these results provide evidence that DNA methylation variability and gene expression variability could be mediated by the sequence-specific binding of transcription factors, such as *ELF1* in neutrophils. Future studies will be required to further investigate the functional relevance of the observed correlation.

## Discussion

In this study, we investigated the transcriptional and epigenetic variability that enables immune cells to rapidly adapt to environmental changes. To this end, we devised a novel analytical strategy to assess inter-individual variability of gene expression and DNA methylation as a measure of functional plasticity across three immune cell types.

A key insight from our integrative analyses is that neutrophils exhibit substantially increased variability of both gene expression and DNA methylation compared to monocytes and T cells (Tables S1 and S6). In neutrophils, genes with important functions in intracellular signaling, cell adhesion, and motility showed increased variability (Tables S2 and S7). Importantly, a subset of these genes were found to be under epigenetic control, such as *RSAD2*, a gene involved in interferon-mediated immune response (Figure 3). Neutrophils play a diverse role in innate immune defense and act as effector cells of adaptive immunity. They use multiple sophisticated molecular mechanisms to locate and contain pathogens, including phagocytosis, degranulation, and NETosis, i.e. the generation of neutrophil extracellular traps, NETs [55,56]. The phenotypic and functional heterogeneity of neutrophils has recently been recognized [57]. Here, we add to these data, providing evidence that increased variability of gene expression and DNA methylation relate to functional diversity and effective adaptability during homeostatic and potentially pathogenic immune processes.

For our analyses we exploited the unique resource provided by the BLUEPRINT Human Variation Panel [24], enabling us to conduct the most comprehensive study of molecular variability in primary cell types to date. It allowed us to perform systematic, paired analyses across cell types and individuals, thus accounting to a large extent for potential differences related to sample processing and donor characteristics. However, we acknowledge that it is not possible to rule out a role for other confounding factors: Heterogeneity may also be partly explained by differing stages and rates of cell activation and cell death during experimental processing, as well as unaccounted environmental effects such as circadian rhythm, diet, physical activity, and psychological stress, which could affect one cell type more than the other(s).

Differences in the proportions of cellular subpopulations may contribute to overall elevated variability between individuals. We have thus assessed the expression profiles of a number of genes that identify distinct cellular subpopulations of neutrophils [57]: *CXCR4*, *CD63*, *CD62L* (also known as *SELL*), and *CD49* (also known as *ITGA4*). We did not observe inter-individual gene expression differences of surface markers corresponding to known neutrophil subpopulations, with the exception of CD49 (Figure S10). We note that *CD49* gene expression levels did not correlate with neutrophil granularity (BH-corrected *P* = 0.89, Spearman’s rank correlation). These data suggest that variation in neutrophil subpopulations is unlikely to be a main determinant of increased inter-individual variability. Future studies are required to corroborate these results and to determine whether uncharacterized cellular subpopulations may contribute to the observed heterogeneity. Novel transcriptome and epigenome profiling techniques investigating gene expression and DNA methylation variability at the level of single cells will provide valuable additional information [58–60].

We have prepared all data sets generated in this study as an easily accessible and freely available online resource, comprising all results that showed statistical significance (n=3,378) [61]. The portal is aimed to enable the research community to mine all results generated by our analyses and to conduct follow-up research into the plasticity of immune cells. In such studies, hypervariable gene-phenotype associations (Tables S3 and S8) can be further characterized using experimental approaches. For example, gene expression and DNA methylation hypervariability could be correlated to pathophysiological triggers of immune responses, such as interferon-γ and lipopolysaccharide [62].

## Conclusions

We found that neutrophils show increased variability in both their gene expression and DNA methylation patterns compared to monocytes and T cells. Our data suggest that increased variability in neutrophils may lead to cellular plasticity, enabling rapid adaptation to new or changing environments such as inflammation and pathogen intrusion. A detailed molecular understanding of the role of cellular heterogeneity in the human immune system is crucial to specifically target a pathogenic cellular subset without compromising immunity, ultimately advancing therapeutic design and treatment strategies in hematopoietic and immunological diseases.

## Methods

### Sample collection and cell isolation

As part of the BLUEPRINT Human Variation Panel, a total of 202 healthy blood donors were recruited from the Cambridge NIHR BioResource [63]. Sample material was obtained at the NHS Blood and Transplant Centre in Cambridge (UK) with informed consent (REC 12/EE/0040). Donors were on average 55 years of age (range, 20^−^75 years) and 46% of donors were male. Blood was processed within three hours of collection. Whole blood was used to purify CD14^+^CD16^−^ monocytes, CD66b^+^CD16^+^ neutrophils, and naïve CD4^+^CD45RA^+^ T cells using a multistep purification strategy. The purity of each cell preparation was assessed by multi-color fluorescence-activated cell sorting (FACS). Purity was on average 95% for monocytes, 98% for neutrophils, and 93% for CD4^+^ T cells. Purified cell aliquots were pelleted, stored at −80°C, and transported to the processing institutes. Further details about the experimental protocols and quality control assessments are provided by the BLUEPRINT Human Variation Panel [24].

### RNA-sequencing assay and data preprocessing

RNA-seq sample preparation and library creation were performed for monocytes and neutrophils at the Max Planck Institute for Molecular Genetics (Germany), and for T cells at McGill University (QC, CA). Purified cell aliquots were lysed and RNA extracted using TRIZOL reagent (Life Technologies) following the manufacturer’s protocol. Sequencing libraries were prepared using a TruSeq Stranded Total RNA Kit with Ribo-Zero Gold (Illumina). Adapter-ligated libraries were amplified and indexed via PCR. Libraries were sequenced using 100 bp single-end reads for monocytes and neutrophils, and paired-end reads for T cells. Reads from each RNA-seq library were assessed for duplication rate and gene coverage using FastQC [64]. Then, PCR and sequencing adapters were trimmed using Trim Galore. Trimmed reads were aligned to the GRCh37 reference genome using STAR [65]. We used GENCODE v15 to define the annotated transcriptome. Read counts of genes and exons were scaled to adjust for differences in total library size using DESeq2 [66]. Then, we applied ComBat [67] to correct for batch effects. An overview of the RNA-seq data quality assessment is provided in Figure S1.

### Quantification of gene expression

Analyses on RNA-seq data were performed on exon-based read counts per gene. We omitted all genes not expressed in at least 50% of all samples in each of the three cell types, leaving only genes that were robustly expressed in all three cell types. In addition, we included only protein-coding genes, resulting in a final set of 11,980 genes. RNA-seq read counts were converted into expression log counts by applying the formula log2(x^+^1).

### Illumina Infinium HumanMethylation450 assay and data preprocessing

For monocytes and neutrophils, cell lysis and DNA extraction were performed at the University of Cambridge (UK), followed by bisulfite conversion and DNA methylation profiling at University College London (UK). T cells were processed at McGill University (QC, CA). DNA methylation levels were measured using Infinium HumanMethylation450 assays (Illumina) according to the manufacturer’s protocol. All 450K array data preprocessing steps were carried out using established analytical methods incorporated in the R package minfi [68]. First, we performed background correction and dye-bias normalization using NOOB [69], followed by normalization between Infinium probe types with SWAN [70]. Next, we filtered out probes based on the following criteria: (1) median detection *P* ≥0.01 in one or more samples; (2) bead count of less than three in at least 5% of samples; (3) mapping to sex chromosomes; (4) ambiguous genomic locations [71]; (5) non-CG probes; and (6) containing SNPs (MAF ≥0.05) within 2 bp of the probed CG. Finally, we adjusted for batch effects using an empirical Bayesian framework [67], as implemented in the ComBat function of the R package SVA [72]. An assessment of the DNA methylation data quality is shown in Figure S2.

### Quantification of DNA methylation

The final data set that passed quality control consisted of 440,905 CpG sites. DNA methylation values were represented as either M-values or β-values. The methylation M-value is the log2-ratio of the intensities of the methylated probe versus the unmethylated probe on the 450K array, while the β-value is the ratio of the methylated probe intensity and the overall intensity. All analyses of DNA methylation data were performed using M-values. Due to their easier interpretability (i.e. 0^−^% DNA methylation), β-values were used for the visualization of DNA methylation data in most figures.

### Analysis of differential variability

To assess differential variability across the three cell types, we applied a combined statistical approach based on DiffVar [73], which is embedded in the framework of limma [74,75]. DiffVar calculates the median absolute deviation from the group mean (MAD) of expression levels of a particular gene, or DNA methylation at a given CpG site, across all individuals for two conditions, e.g. two distinct cell types. Then, a moderated t-test is used to test for a significant increase or decrease in MAD-value between the two conditions. However, we found that the MAD variability measurement employed by DiffVar is correlated with mean levels (Figures S3 and S7), which could potentially confound the assessment of variability. Therefore, we included an additional measurement of variability that corrects for the dependency of variability measurements on the mean [8], here referred to as EV (gene expression variability value) and MV (DNA methylation variability value). The corresponding algorithm models variance as a function of the mean, and then calculates the ratio of the observed variance to expected variance in order to get a variability measurement independent of the mean. Differential variability was tested in three group-wise comparisons. Statistical significance was defined as BH-corrected [76] *P* <0.05 and EV/MV difference ≥10% relative to the observed range of EV-/MV-values. For each cell type, both contrasts in which the cell type is involved were considered to define statistically significant differential variability. For example, for a gene to be a neutrophil-specific HVG, it must show significantly increased variability in both the comparison versus monocytes and versus T cells. For a gene to be classified as hypervariable across two cell types (shared hypervariability), it must exhibit significantly increased variability in the two corresponding cell types, but low variability in the third. Thus, no gene can appear in more than one list. The statistical tests were performed in a paired fashion, taking into account that all three cell types were derived from the same individuals. This procedure corrects for potential differences related to individuals and sample processing.

### Analysis of variability common to all three cell types

To identify HVGs common to all three cell types, we applied a rank-based approach. We ordered both MAD- and EV-values of all genes in the three cell types from high to low variability, and then took the top n genes with highest variability across all three cell types, where n corresponds to the mean number of results obtained for the gene lists of differential variability. Specifically, n=271 for gene expression variability, and n=212 for DNA methylation variability.

### Gene set enrichment analyses

For HVGs, we applied GOseq using the default parameters, and set ‘use_genes_without_cat’ = FALSE, thus ignoring genes without an annotated category for the calculation of *P*-values [28]. With regards to HVPs, we analyzed the biological functions of flanking genes with GREAT [43] using the standard parameters: association rule = basal ^+^ extension (constitutive 5 kb upstream, kb downstream, up to 1 Mb extension); curated regulatory domains = included. In both analyses we used the set of analyzed features as background, and the cutoff for statistical significance was set at BH-corrected *P* <0.25.

### Gene co-expression network and pathway analysis

For neutrophil-specific HVGs not associated with *cis* genetic variants in the BLUEPRINT Human Variation Panel, we first constructed a co-regulation network by calculating gene expression correlations. The threshold of gene correlations was set at Pearson’s r >0.6. Unconnected genes were removed. The resulting correlation network was then further analyzed using Cytoscape [77]. Clusters were identified by the agglomerative clustering method FAG-EC [78] of the ClusterViz plugin. Enrichment analyses of resulting gene clusters were performed using clueGO [79], setting the Kappa score to 0.4 and the cutoff for statistical significance at BH-corrected *P* <0.05. All networks were visualized using Gephi [80].

### Correlation analyses

Associations between both gene expression and DNA methylation levels with donor-specific quantitative traits, cellular parameters, as well as weather and seasonal effects were assessed by calculating Spearman’s rank correlation coefficients (rho) and their corresponding *P*-values. Results were considered statistically significant at BH-corrected *P* <0.05. This threshold was also used for the correlation analyses between DNA methylation and gene expression data.

### Analyses of seasonal effects

We downloaded historical raw weather data for the minimum and maximum daily temperature in London Heathrow (UK) for the period of data collection from the National Climatic Data Centre (USA) [81]. We applied linear interpolation to account for missing values. Additionally, we downloaded daylight hours for London [82]. The obtained data were then correlated with gene expression and DNA methylation values corresponding to the date of blood donation using Spearman’s rank correlation coefficient (see details above).

### Analyses of sex-specific differential gene expression

In each cell type, mean gene expression and DNA methylation differences between male and female donors were identified using limma [74,75]. A moderated t-test was performed and statistical significance defined as BH-corrected *P* <0.05 and log-fold change ≥1. Results could be driven by differences in menopause status between female donors. Therefore, we performed the same analysis on only the subset of donors whom are younger than 50 years, and obtained very similar results compared to the complete donor group.

### Functional annotation of hypervariable CpGs

For the enrichment analyses with regards to gene elements and epigenomic features, we used the annotation provided by the Illumina 450K array manifest. Enrichment was assessed by repeated random sampling (n=1,000) using all probes that passed quality control (n=440,905).

### Transcription factor motifs analysis at gene promoter regions

Consensus transcription factor binding motifs were retrieved from the database ‘JASPAR_CORE_2016 _vertebrates.meme’ [83]. Using FIMO [84], we scanned for transcription factor binding motifs (*P* < 1×10-5) at promoter regions, defined as ±500 bp around the transcription start site of genes listed in the reference gene set ‘UCSC.hg19.knownGene’.

#### Programming language

If not indicated otherwise, analyses were performed using R v3 (R Development Core Team, 2008) and Bioconductor [85].

## List of Abbreviations

HIV: human immunodeficiency virus
RNA-seq: RNA sequencing
HVG: hypervariable gene
HVP: hypervariable
CpG: position
FACS: fluorescence-activated cell sorting
MAD: median absolute deviation
EV: gene expression variability value
MV: DNA methylation variability value

## Declarations

### Ethics approval and consent to participate

All sample material was obtained at the NHS Blood and Transplant Centre in Cambridge (UK) with informed consent (REC 12/EE/0040).

### Consent for publication

Not applicable.

## Authors’ Contributions

S. Ecker and D.S. Paul designed the study. S. Ecker, D.S. Paul, L. Chen, V. Pancaldi, F.O. Bagger, E. Carrillo de Santa Pau, D. Juan, N. Sidiropoulos, N. Rapin, and A. Merkel analyzed data. All other authors provided samples or analytical tools. D.S. Paul and S. Ecker wrote the manuscript. D. Rico, A. Valencia, S. Beck, N. Soranzo, and D.S. Paul supervised the study. All authors read and approved the final manuscript.

## Acknowledgments

We would like to thank K. Pearce and M. Kristiansen (UCL Genomics) for processing the Illumina Infinium HumanMethylation450 BeadChips; D. Balzereit, S. Dökel, A. Kovacsovics, and M. Linser (Max Planck Institute for Molecular Genetics) for help with generating the RNA-seq data; B. Phipson (Murdoch Childrens Research Institute) and H.C. Bravo (University of Maryland) for advice on statistical analyses; C. Bock (CeMM Research Center for Molecular Medicine of the Austrian Academy of Sciences) for useful discussions; A. Orozco (University of Costa Rica) for technical support; V. Naranbhai, B. Fairfax, and J. Knight (University of Oxford) for providing access to the neutrophil gene expression data set for replication; and L. Phipps for proofreading the manuscript. We gratefully acknowledge the participation of all NIHR Cambridge BioResource volunteers, and thank the Cambridge BioResource staff for their help with volunteer recruitment. We thank members of the Cambridge BioResource SAB and Management Committee for their support of our study and the NIHR Cambridge Biomedical Research Centre for funding. S. Ecker is supported by a “la Caixa” pre-doctoral fellowship. V. Pancaldi is supported by a FEBS Long-Term Fellowship. F.O. Bagger is supported by The Lundbeck Foundation. K. Downes is funded as a HSST trainee by NHS Health Education England. M. Frontini is supported by the BHF Cambridge Centre of Excellence (RE/13/6/30180). S. Beck acknowledges support from the Wellcome Trust (WT99148), a Royal Society Wolfson Research Merit Award (WM100023), and the EU-FP7 Project EPIGENESYS (257082). N. Soranzo’s research is supported by the Wellcome Trust (WT098051 and WT091310), EU-FP7 Project EPIGENESYS (257082), and NIHR BRC.

BLUEPRINT Consortium: Cornelis A. Albers (Radboud University), Vyacheslav Amstislavskiy (Max Planck Institute for Molecular Genetics), Sofie Ashford (University of Cambridge), Lorenzo Bomba (Wellcome Trust Sanger Institute), David Bujold (McGill University), Frances Burden (University of Cambridge), Stephan Busche (McGill University), Maxime Caron (McGill University), Shu-Huang Chen (McGill University), Warren A. Cheung (McGill University), Laura Clarke (European Bioinformatics Institute), Irina Colgiu (Wellcome Trust Sanger Institute), Avik Datta (European Bioinformatics Institute), Oliver Delaneau (University of Geneva), Heather Elding (Wellcome Trust Sanger Institute), Samantha Farrow (University of Cambridge), Diego Garrido-Martín (Centre for Genomic Regulation), Bing Ge (McGill University), Roderic Guigo (Centre for Genomic Regulation), Valentina Iotchkova (European Bioinformatics Institute), Kousik Kundu (Wellcome Trust Sanger Institute), Tony Kwan (McGill University), John J. Lambourne (University of Cambridge), Ernesto Lowy (European Bioinformatics Institute), Daniel Mead (Wellcome Trust Sanger Institute), Farzin Pourfarzad (Sanquin Research and Landsteiner Laboratory), Adriana Redensek (McGill University), Karola Rehnstrom (University of Cambridge), Augusto Rendon (University of Cambridge), David Richardson (European Bioinformatics Institute), Thomas Risch (Max Planck Institute for Molecular Genetics), Sophia Rowlston (University of Cambridge), Xiaojian Shao (McGill University), Marie-Michelle Simon (McGill University), Marc Sultan (Max Planck Institute for Molecular Genetics), Klaudia Walter (Wellcome Trust Sanger Institute), Steven P. Wilder (European Bioinformatics Institute), Ying Yan (Wellcome Trust Sanger Institute), Stylianos E. Antonarakis (University of Geneva), Guillaume Bourque (McGill University), Emmanouil T. Dermitzakis (University of Geneva), Paul Flicek (European Bioinformatics Institute), Hans Lehrach (Max Planck Institute for Molecular Genetics), Joost H. A. Martens (Radboud University), Marie-Laure Yaspo (Max Planck Institute for Molecular Genetics), Willem H. Ouwehand (University of Cambridge).

